# Discrete Cognitive Resolution in Alzheimer’s Disease: Cross-Cohort Reanalysis of ADNI and NACC Longitudinal Data

**DOI:** 10.64898/2026.06.18.733215

**Authors:** Alia Wu

**Affiliations:** Redline Rising, ISNI: 0000 0005 2969 3119, New York, NY, USA

**Author notes:** Corresponding author: Alia Wu, Redline Rising.

**Keywords:** Alzheimer’s disease, dementia, finite-state model, cognitive diagnostic model, ADAS-Cog, ADNI, NACC

## Abstract

**INTRODUCTION:** Do continuous cognitive totals capture patient-relevant transitions, or does decline have discrete structure?

**METHODS:** We formalize a discrete cognitive resolution (DCR) model in which decline is loss of binary discriminative coordinates, with deterministic emissions tying items to a shared low-dimensional mask. Using natively discrete item-level data (ADAS-Cog, 688 ADNI participants; MoCA sub-items, 13,323 NACC participants), we tested pre-specified signatures by out-of-sample log-loss against continuous-drift, per-item, and mixed-effects IRT competitors, with a coordinate-label permutation null (S2).

**RESULTS:** DCR beat both pre-specified baselines (ADNI 0.436 vs 0.965; NACC 0.566 vs 0.602). S2 was decisive in ADNI (AUC 0.782; null 0.529, *P <* .001); in NACC the signal concentrated in orientation (AUC 0.718). Mixed-effects IRT achieved lower log-loss than DCR.

**DISCUSSION:** Cognitive decline shows discrete coordinate structure when items are single-coordinate probes. The claim is structural, not predictive; encoding is decisive.

## 1. Background

The April 2026 Cochrane review of amyloid *β* (A*β*)-targeting monoclonal antibodies for mild cognitive impairment (MCI) or mild dementia due to Alzheimer’s disease (AD) synthesized 17 randomized trials (20,342 participants) and concluded that average effects on cognitive decline and dementia severity at 18 months were absent or trivial, while amyloid-related imaging abnormalities were increased [1]. The finding should be interpreted carefully: phase 3 trials of lecanemab and donanemab reported statistically significant slowing of decline in selected early populations [2,3], the pooling of mechanistically heterogeneous agents has drawn sustained criticism, and combining older and newer antibodies in one meta-analysis remains contested. Nevertheless the review sharpens a measurement question that is independent of the biology: do standard continuous outcomes capture patient-relevant transitions in cognitive capacity? If a patient’s ability to navigate, manage medications, or follow conversations depends on specific discriminative capacities rather than a scalar total, then the distinction between which capacities are lost and how many points have been lost is clinically material.

Most trial and cohort analyses summarize cognition through continuous or ordinal totals— the Mini-Mental State Examination, Montreal Cognitive Assessment (MoCA), Clinical Dementia Rating Sum of Boxes (CDR-SB), or the Alzheimer’s Disease Assessment Scale–Cognitive subscale (ADAS-Cog) [4–7]. These instruments are indispensable for staging, regulation, and comparison. Their convenience can conceal a stronger assumption: that decline is well represented as smooth change in a scalar score. Alzheimer’s disease and related dementias show heterogeneity, nonlinear acceleration, fluctuations, domain-specific vulnerability, and individual trajectory differences [8– 10]; when assessments occur every 6 to 12 months, a sequence of latent transitions can appear as a continuous average slope. Mature latent-state and change-point models already encode the possibility of discrete transitions [11,12], and cognitive diagnostic models classify examinees into latent skill profiles from structured assessment data [13–15], but these frameworks are rarely applied to longitudinal item-level dementia data—the setting where discrete coordinate loss would be most directly observable.

This paper develops a complementary modeling layer and tests whether its pre-specified signatures are present in two independent cohorts: item-level ADAS-Cog data from 688 ADNI participants and MoCA sub-item data from 13,323 NACC participants. We formalize a discrete cognitive resolution (DCR) model in which decline is loss of binary discriminative coordinates in a finite register, state the constraint that distinguishes it from a generic hidden Markov model—that emissions are deterministic projections through a shared low-dimensional mask—name mask identifiability as the central open problem, and test whether five pre-specified diagnostic signatures replicate across cohorts, instruments, and scales. A key methodological result is that encoding matters: the model requires natively discrete item-level data with single-coordinate loading, and applying it to continuously-scored test-level aggregates produces spurious failures attributable to the encoding, not the model. The contribution is structural, not predictive: we show that cognitive decline has discrete coordinate organization, a claim that no continuous model can produce regardless of parameterization.

## 2. Methods

### 2.1 The discrete cognitive resolution model

A cognitive task environment is represented by *d* binary discriminative coordinates. At time *t* an individual has an active resolving mask *m*(*t*) ∈ *{*0, 1*}*^*d*^, where *m*_*j*_(*t*) = 1 means coordinate *j* supports reliable discrimination. Resolving power is *ρ*(*t*) = ∑_*j*_ *m*_*j*_(*t*). A stimulus or task demand is a vector *x* ∈ *{*0, 1*}*^*d*^; the effective record is the projection *R*_*m*(*t*)_(*x*) = *m*(*t*) ⊙ *x*. Two states are indistinguishable to the individual when *m*(*t*) ⊙ *x* = *m*(*t*) ⊙ *y*, so active coordinates partition the environment into 2^*ρ*^ equivalence classes, each collapsing 2^*d*−*ρ*^ states distinguishable to a higher-resolution observer. A coordinate-loss event *m*_*j*_ : 1 → 0 halves the distinguishable classes within the engaged subsystem, from 2^*ρ*^ to 2^*ρ*−1^ (full derivation, Supplement S1). The model does not assert literal neural bits; it asserts that clinical tasks decompose into finite discriminative dimensions and that loss of a dimension produces structured coarse-graining rather than uniform noise. The discrete layer concerns the functional consequence once a coordinate falls below the threshold for reliable discrimination, and is compatible with continuous neurobiology: synaptic, network, tau, vascular, inflammatory, sleep, sensory, and medication factors all change the probability that a coordinate remains active, but the model’s predictions concern the observable consequences of the transition, not the mechanism. In this respect the model is analogous to phase transitions in physics, where continuous changes in temperature produce discrete changes in observable state; the claim is not that neural tissue is digital but that the mapping from neural capacity to behavioral discrimination has thresholds.

The model’s identifying claim is that emissions are deterministic projections through a shared mask: for task *k* requiring coordinate set *τ*_*k*_,

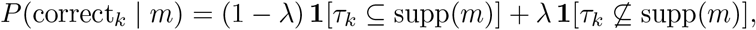

with lapse rate *λ*. The emission distribution across all tasks is generated by the single *d*-dimensional mask, not by free per-task parameters. A generic hidden Markov model fits each state’s emission vector freely, costing *O*(2^*d*^) parameters across the state lattice; the DCR constraint ties emissions through the mask at *O*(*d*). This restriction is the model’s identifying content and the source of its statistical leverage: it predicts that failures across unrelated tasks at a given time are organized by a common set of lost coordinates rather than being task-specific. That cross-task coordinate consistency is the signature that separates DCR from spreading-activation or item-specific confusability accounts, which predict task-local error structure. A spreading-activation model, for instance, would predict that errors propagate through associative networks regardless of coordinate membership; DCR predicts that errors cluster by coordinate.

Every clinical use of the model routes through the inferred mask *^* (*t*). With *d* coordinates there are 2^*d*^ possible masks, and from accuracy data alone one cannot in general separate “coordinate inactive” from “coordinate active but failing stochastically” without an externally fixed coordinate dictionary mapping each task to the coordinates it requires. We treat the dictionary as a preregistered modeling input and treat mask identifiability as the primary feasibility question.

### 2.2 Pre-specified diagnostic signatures

Five signatures were specified before examining cohort-level results.

**S1 (model comparison):** A DCR model should achieve lower out-of-sample item-level logloss than a continuous-drift alternative (an IRT-style logistic with per-patient ability linear in time and shared per-task difficulty [16]) and a per-item baseline (independent logistic per item with no pooling—a flexible baseline that ignores shared structure). All models are trained on the first two-thirds of each patient’s visits and evaluated on the held-out final third.

**S2 (equivalence-class errors):** Item failure should be predicted by whether the item’s coordinate is currently weak, measured leave-one-item-out on other items sharing that coordinate. Because items share a global severity gradient—in advanced disease most items fail together regardless of coordinate—a raw AUC above chance is insufficient evidence of coordinate-specific organization. We therefore pre-specified three companion analyses: (i) a permutation null that shuffles coordinate labels across items while preserving the number of items per coordinate, recomputing the AUC 1000 times, so that the null absorbs whatever co-failure any same-sized grouping would produce; (ii) a per-coordinate decomposition of the AUC, identifying which coordinates carry the signal; and (iii) within- versus between-coordinate pairwise item agreement, a descriptive check that items sharing a coordinate co-fail more than items on different coordinates.

**S3 (ordered vulnerability):** Tasks requiring more coordinates should fail earlier, because the probability that all required coordinates remain active decreases with the number required. This prediction holds under equal per-coordinate hazard but may be confounded by domain-specific vulnerability. **S4 (interaction-order collapse):** The Walsh-Hadamard decomposition of performance should show loss of high-order interaction terms before low-order terms. Both S3 and S4 require multi-coordinate items and are not testable on single-coordinate batteries.

**S5 (mask vs. total):** If the mask captures clinically relevant structure beyond overall severity, it should predict functional outcomes better than the total score. The honest bar: the mask must beat a fair saturated item-level baseline, not merely the total.

### 2.3 Cohorts, encoding, and analysis

Both cohorts are existing longitudinal observational studies analyzed under their Data Use Agreements; no patient-level data reside in the analysis repository. Participants were eligible with ≥ 3 item-level visits.

**ADNI**. In the Alzheimer’s Disease Neuroimaging Initiative [17], item-level ADAS-Cog responses (Q1–Q12, with Q9 excluded for near-zero variance) were obtained from the cognitive behavior item-level export. Sub-item responses (per-word recall across 3 trials, per-command, per-naming-item, per-orientation-item) were aggregated to question-level error scores and binarized using natural pass/fail thresholds for each question (e.g., Q1 word recall: pass if ≤ 3 errors out of 10; Q7 orientation: pass if ≤ 1 error out of 8). No z-score normalization or reference-group computation was applied—each item’s pass/fail criterion reflects the natural scoring structure. This yielded 688 participants (median 5 visits, IQR 4–5, ~6-month intervals; 26,022 item-visit observations). Each item mapped to one of five coordinates following the instrument’s published subscale structure [7]: Q1/Q4/Q8→memory; Q2/Q5/Q10/Q11/Q12→language; Q3→visuospatial; Q6→executive; Q7→orientation.

**NACC**. In the National Alzheimer’s Coordinating Center Uniform Data Set (UDS3) [18], 19 MoCA sub-items (Form C2), each natively scored as binary (0/1) or small ordinal (0–2, 0– 3, 0–5), were binarized at natural pass/fail thresholds (e.g., serial 7 subtraction: pass if all 3 correct; delayed recall: pass if ≥ 3 of 5 words recalled without cue). No z-score normalization was applied. Each sub-item mapped to one of six coordinates following MoCA’s published domain structure [5]: visuospatial (cube copy, clock contour, clock numbers), executive (trail making, clock hands, abstraction), language (naming, sentence repetition, verbal fluency), attention (digit span, letter tapping, serial 7s), memory (delayed recall), and orientation (date, month, year, day, place, city). This yielded 13,323 participants with annual visits and approximately 1,089,005 item-visit observations—an order of magnitude larger than ADNI, providing a stringent independent replication test on a different instrument with different cadence and population characteristics.

Single-coordinate loading ensures the product emission reduces to a single active-probability per item, avoiding the entanglement between coordinates that arises when a single item’s success probability depends on the product of multiple coordinate-specific terms. Coordinate dictionaries (pre-registered Q-matrices) are in Supplement S2; cohort characteristics are in Table 2. The minimum-visit criterion selected a younger, less impaired analytic sample than the full UDS source population (mean age 68.9 vs 71.0 years; baseline CDR-SB 0.79 vs 2.43; 63% vs 42% cognitively normal and 11% vs 31% with dementia), as expected when requiring repeated follow-up favors slower-declining participants (Table 2). Findings should be generalized with this selection in mind.

**Encoding rationale**. An earlier NACC analysis using 12 test-level summary scores (MoCA total, Craft Story, MINT naming, digit span, category fluency, Trail Making, Benson figure) with z-score binarization and multi-coordinate loading produced a reversal: DCR lost to continuous (Δ = −0.147). Investigation revealed this was an encoding artifact: tests like MoCA total load on 5 coordinates, and the product emission *P* (pass) = II_*c*_ *p*_*c*_ drives predictions toward certain failure even when individual coordinates are only mildly impaired (e.g., 0.9^5^ = 0.59). The fix was not to change the model but to use the correct data level: natively discrete sub-items, each loading on one coordinate. The reversal was entirely attributable to encoding.

**Analysis**. Three models were fit per patient on the first two-thirds of visits and evaluated by out-of-sample log-loss on the final third, with patient-level bootstrap 95% confidence intervals (1000 resamples) for the DCR–continuous difference: (1) a continuous-drift IRT-style model with per-patient ability linear in time and shared per-task difficulty, (2) the DCR shared-mask projection model, and (3) a per-item logistic baseline with no pooling across items. A mixed-effects IRT competitor—a per-patient random intercept and slope model (2*n* + *k* effective parameters) with item difficulties pooled across training visits and shrunk toward population values by an empiricalBayes *L*_2_ prior [16]—was added as a strong calibration benchmark. The regularization strength for the continuous competitor (*C* = 2.0) and per-item baseline (*C* = 5.0) were set to reasonable defaults and not tuned on the test set. For the S5 signature, the CDR Sum of Boxes (CDR-SB) at the last available visit was used as the prospective functional outcome, predicted from training-window data using the inferred mask, the total score, and the saturated item vector [6]. A pre-specified sensitivity analysis varied *λ* ∈ *{*0.02, 0.05, 0.10*}*, train fraction *{*0.50, 0.66, 0.75*}*, and the NACC recall threshold (Supplement S3).

## 3. Results

### DCR is competitive with the pre-specified baselines (S1)

DCR achieved out-of-sample item-level log-loss of 0.436 in ADNI versus 0.965 (continuous) and 0.833 (per-item), a large and significant advantage (Δ = 0.529; bootstrap 95% CI [0.44, 0.63]; Table 1, Figure 1A). In NACC the pattern replicated in an independent cohort 19 times larger, with a different instrument and annual cadence: 0.566 versus 0.602 and 0.800 (Δ 95% CI [0.029, 0.043]; Figure 2A). The DCR advantage (Δ = 0.529) is large in ADNI, indicating that real ADAS-Cog trajectories contain coordinate structure that smooth-drift models miss; the NACC advantage (Δ = 0.036) is significant but smaller. Two factors likely contribute to the difference: (1) ADAS-Cog items span a wider difficulty range and were designed to detect impairment across the MCI-to-dementia spectrum, whereas many MoCA sub-items show ceiling effects in cognitively-normal populations; (2) ADNI’s semi-annual assessments provide denser temporal sampling than NACC’s annual visits, giving the model more training data per patient and more opportunity to detect coordinate transitions within the training window.

**Table 1.**
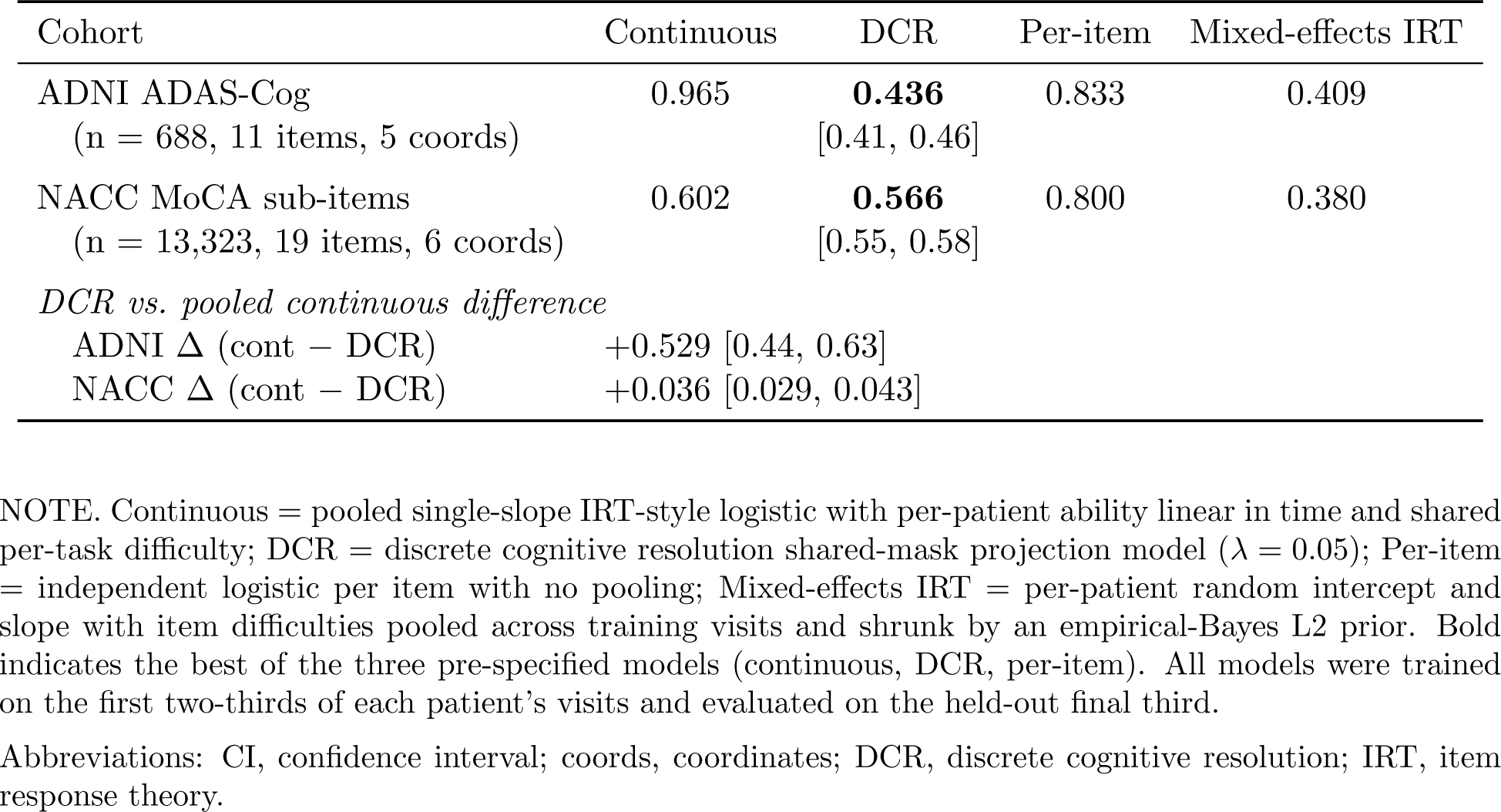
Out-of-sample item-level log-loss (lower is better) for competing models on two independent cohorts. Both cohorts use natively discrete items with single-coordinate loading. The mixed-effects IRT model fits per-patient random intercepts and slopes (2n+k effective parameters); it achieves the lowest log-loss but has no concept of coordinates and cannot produce the S2 signature. Bootstrap 95% CIs (1000 patient-level resamples) are shown for DCR.

| Cohort | Continuous | DCR | Per-item | Mixed-effects IRT |
| --- | --- | --- | --- | --- |
| ADNI ADAS-Cog<br>(n = 688, 11 items, 5 coords) | 0.965 | **0.436**<br>[0.41, 0.46] | 0.833 | 0.409 |
| NACC MoCA sub-items<br>(n = 13,323, 19 items, 6 coords) | 0.602 | **0.566**<br>[0.55, 0.58] | 0.800 | 0.380 |
| *DCR vs. pooled continuous difference* |  |  |  |  |
| ADNI $\Delta$ (cont – DCR) | +0.529 [0.44, 0.63] | | | |
| NACC $\Delta$ (cont – DCR) | +0.036 [0.029, 0.043] | | | |

NOTE. Continuous = pooled single-slope IRT-style logistic with per-patient ability linear in time and shared per-task difficulty; DCR = discrete cognitive resolution shared-mask projection model ( $\lambda = 0.05$ ); Per-item = independent logistic per item with no pooling; Mixed-effects IRT = per-patient random intercept and slope with item difficulties pooled across training visits and shrunk by an empirical-Bayes L2 prior. Bold indicates the best of the three pre-specified models (continuous, DCR, per-item). All models were trained on the first two-thirds of each patient’s visits and evaluated on the held-out final third.
Abbreviations: CI, confidence interval; coords, coordinates; DCR, discrete cognitive resolution; IRT, item response theory.

**Table 2.**
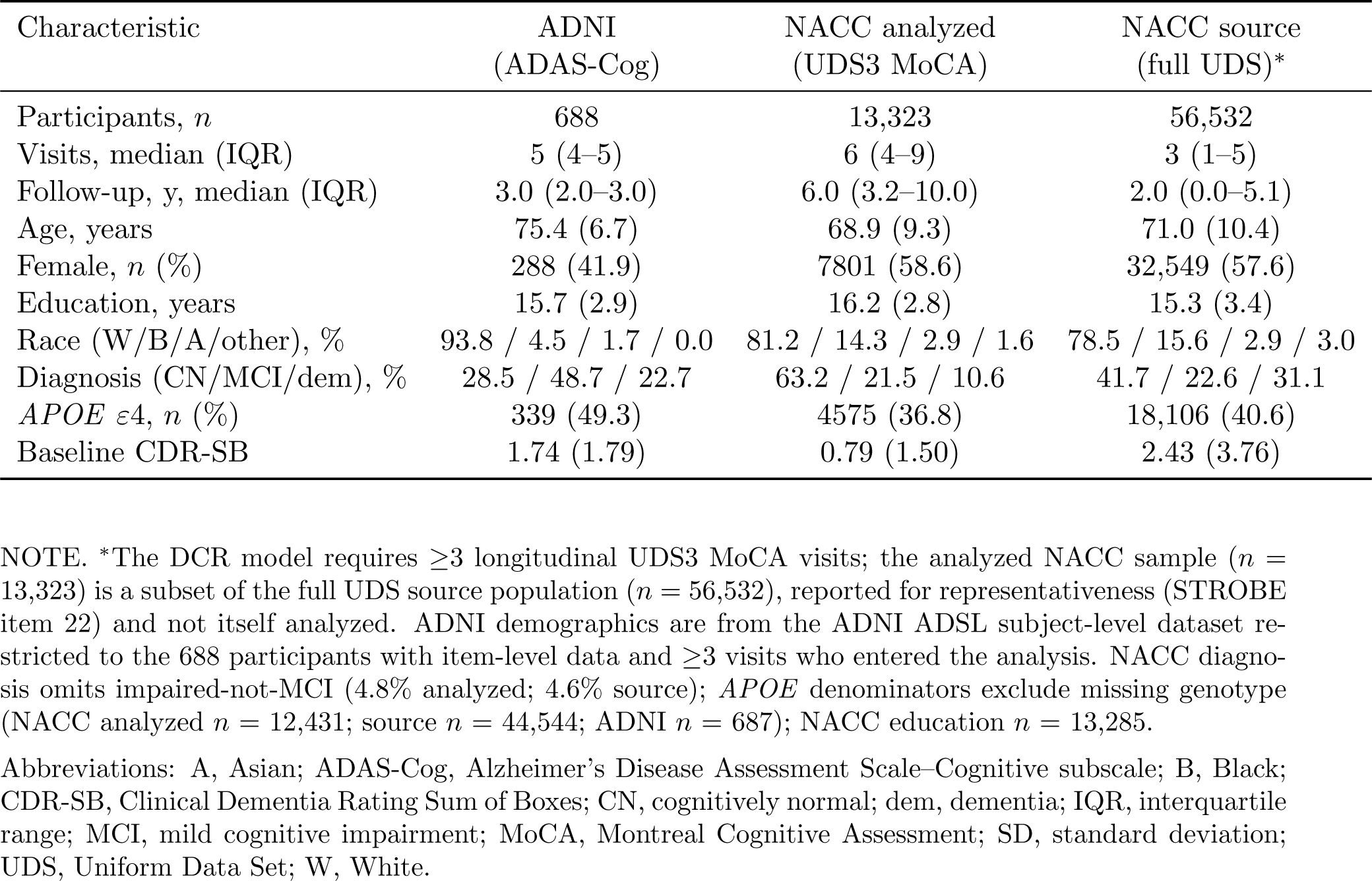
Cohort characteristics at baseline for the analytic sample (participants entering the analysis). Values are mean (SD) unless noted. NACC values generated by the released analysis pipeline restricted to the analyzed participants with missing/unknown codes cleaned; ADNI values from the ADNI ADSL subject-level dataset restricted to the item-level population.

| Characteristic | ADNI<br>(ADAS-Cog) | NACC analyzed<br>(UDS3 MoCA) | NACC source<br>(full UDS)* |
| --- | --- | --- | --- |
| Participants, *n* | 688 | 13,323 | 56,532 |
| Visits, median (IQR) | 5 (4–5) | 6 (4–9) | 3 (1–5) |
| Follow-up, y, median (IQR) | 3.0 (2.0–3.0) | 6.0 (3.2–10.0) | 2.0 (0.0–5.1) |
| Age, years | 75.4 (6.7) | 68.9 (9.3) | 71.0 (10.4) |
| Female, *n* (%) | 288 (41.9) | 7801 (58.6) | 32,549 (57.6) |
| Education, years | 15.7 (2.9) | 16.2 (2.8) | 15.3 (3.4) |
| Race (W/B/A/other), % | 93.8 / 4.5 / 1.7 / 0.0 | 81.2 / 14.3 / 2.9 / 1.6 | 78.5 / 15.6 / 2.9 / 3.0 |
| Diagnosis (CN/MCI/dem), % | 28.5 / 48.7 / 22.7 | 63.2 / 21.5 / 10.6 | 41.7 / 22.6 / 31.1 |
| *APOE* ε4, *n* (%) | 339 (49.3) | 4575 (36.8) | 18,106 (40.6) |
| Baseline CDR-SB | 1.74 (1.79) | 0.79 (1.50) | 2.43 (3.76) |

NOTE. \*The DCR model requires $\geq 3$ longitudinal UDS3 MoCA visits; the analyzed NACC sample ( $n = 13,323$ ) is a subset of the full UDS source population ( $n = 56,532$ ), reported for representativeness (STROBE item 22) and not itself analyzed. ADNI demographics are from the ADNI ADSL subject-level dataset restricted to the 688 participants with item-level data and $\geq 3$ visits who entered the analysis. NACC diagnosis omits impaired-not-MCI (4.8% analyzed; 4.6% source); *APOE* denominators exclude missing genotype (NACC analyzed $n = 12,431$ ; source $n = 44,544$ ; ADNI $n = 687$ ); NACC education $n = 13,285$ .
Abbreviations: A, Asian; ADAS-Cog, Alzheimer’s Disease Assessment Scale–Cognitive subscale; B, Black; CDR-SB, Clinical Dementia Rating Sum of Boxes; CN, cognitively normal; dem, dementia; IQR, interquartile range; MCI, mild cognitive impairment; MoCA, Montreal Cognitive Assessment; SD, standard deviation; UDS, Uniform Data Set; W, White.

**Figure 1.**
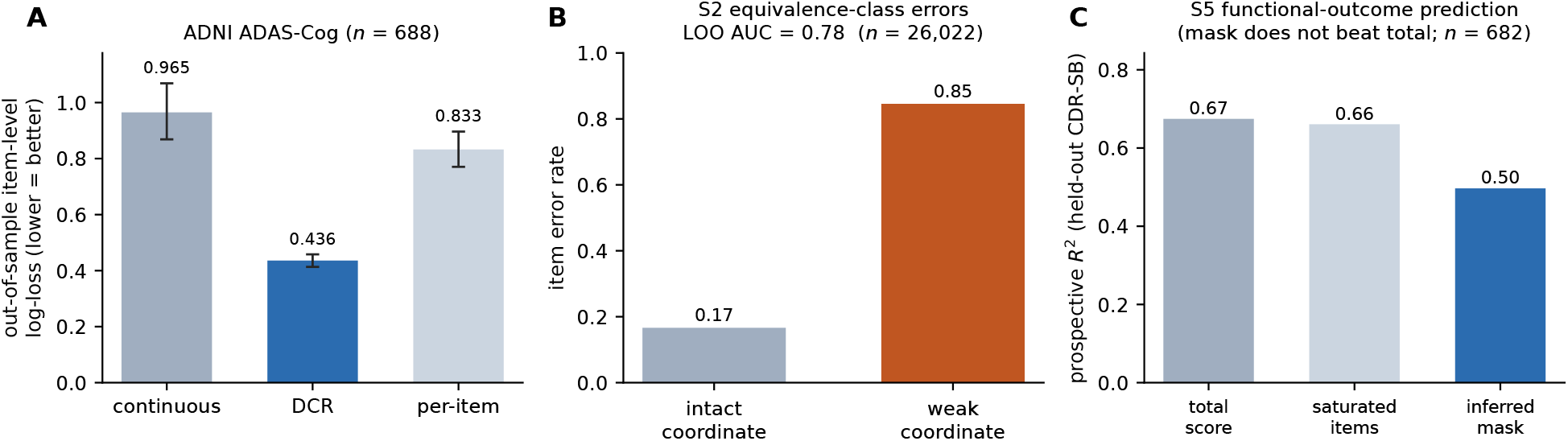
ADNI reanalysis (*n* = 688, ADAS-Cog items). (A) Out-of-sample item-level log-loss for the continuous-drift, DCR, and per-item models; error bars are patient-level bootstrap 95% CIs (1000 resamples). (B) S2 equivalence-class error structure: item error rate when the item’s coordinate is intact versus weak, leave-one-item-out (AUC = 0.78; *n* = 26,022 item-visits). (C) S5: prospective held-out *R*^2^ for CDR-SB from the total score, saturated item vector, and inferred mask.

**Figure 2.**
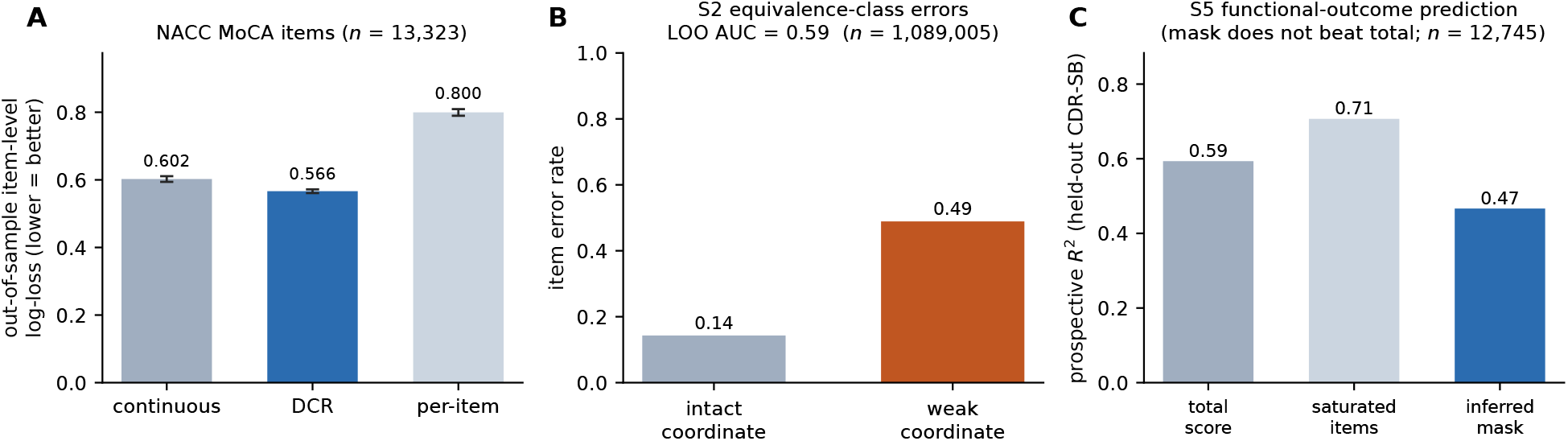
NACC reanalysis (*n* = 13,323, MoCA sub-items), parallel to Figure 1. (A) Model comparison with patient-level bootstrap 95% CIs. (B) S2 equivalence-class error structure, leave-one-item-out (AUC = 0.59; *n* = 1,089,005 item-visits). (C) S5 prospective held-out *R*^2^ for CDR-SB (*n* = 12,745; mask does not beat total).

### Equivalence-class error structure is decisive in ADNI (S2)

Using leave-one-item-out estimation of coordinate weakness (excluding the target item to prevent circularity), item failure was predicted by coordinate status with AUC = 0.782 in ADNI (*n* = 26,022 item-visits; error rate 0.85 on weak-coordinate items vs 0.17 intact). This is decisively coordinate-specific rather than an artifact of global co-decline: against the pre-specified permutation null that shuffles coordinate labels, the observed AUC lies ≈ 5 standard deviations above the null mean (0.529 *±* 0.048; 95% null interval [0.46, 0.66]; *P <* .001, 1000 permutations). Both multi-item coordinates contribute (per-coordinate AUC 0.673 memory, 0.614 language; the visuospatial, executive, and orientation coordinates are each represented by a single ADAS item and are excluded from S2 for lack of cross-task evidence). Pairwise item agreement is far higher within coordinates than between them (85.3% vs 51.2%; excess 34.1 percentage points). This is the defining DCR signature: failures across unrelated tasks are organized by a common set of lost coordinates rather than being task-specific. **S2 concentrates in orientation in NACC**. In NACC the aggregate AUC was 0.593 (*n* = 1,089,005 item-visits; error rate 0.49 vs 0.14) but only marginally above its permutation null (0.576 *±* 0.013; *P* = .093): because MoCA sub-items co-decline strongly and many sit near ceiling, even random groupings reproduce much of the apparent consistency (the NACC null, 0.576, is well above the ADNI null, 0.529). The per-coordinate decomposition reveals marked heterogeneity (AUC 0.718 orientation, 0.599 executive, 0.567 language, 0.547 attention, 0.514 visuospatial; memory excluded as single-item coordinate). The structure concentrates exactly where the model predicts it should be visible—orientation, the coordinate with the widest discriminative range and least ceiling compression (6 items; within-coordinate agreement 91.6% vs 73.5% between)—while ceiling-prone coordinates dilute the pooled AUC. The direction matches ADNI; the attenuated aggregate reflects MoCA ceiling effects and annual cadence, not an absence of coordinate organization.

### A mixed-effects IRT model wins on log-loss in both cohorts

A mixed-effects IRT competitor achieved lower log-loss than DCR on both cohorts (0.409 vs 0.436 ADNI; 0.380 vs 0.566 NACC; Table 1). This is expected: for *n* patients and *k* items it fits 2*n* + *k* effective parameters against DCR’s *d* coordinate trajectories with a shared deterministic emission, and logloss is the metric per-patient calibration optimizes. It does not adjudicate coordinate structure— the mixed model has no concept of coordinates and cannot produce the S2 signature regardless of parameterization. The NACC gap is substantially larger, consistent with MoCA ceiling effects and annual cadence limiting DCR’s leverage there. Change-point and Gaussian-process trajectory models are the natural further comparators at higher sampling frequencies.

### Functional-outcome prediction is a deliberate null (S5)

In prospective held-out prediction of CDR-SB at the final visit, the total score (*R*^2^ = 0.674 ADNI, 0.593 NACC) and the saturated item vector (0.661 ADNI, 0.707 NACC) both exceeded the inferred mask (0.497 ADNI, 0.467 NACC; Figures 1C, 2C). The mask does not add incremental value over the total for a global severity outcome that the total already captures efficiently. This is expected: CDR-SB depends primarily on overall severity, which the total score captures efficiently; the mask’s *d*-dimensional compression is too coarse to compete with the total for a global outcome and too coarse to capture the item-level detail that the saturated vector exploits. In NACC the 19-item saturated vector substantially exceeds the total (*R*^2^ 0.707 vs 0.593), showing that item-level detail carries clinically relevant information the total discards—but the mask’s low-dimensional compression does not capture this advantage. The mask’s potential value lies in predicting domain-specific functional outcomes (e.g., navigation impairment from visuospatial loss, medication errors from executive loss), which CDR-SB does not provide. We report S5 as a deliberate, honest null on incremental value for a global outcome, not as evidence against the model.

## 4. Discussion

The DCR model’s structural signature replicates across two independent cohorts, instruments, and cadences. Cross-task coordinate consistency (S2 AUC 0.782 ADNI, 0.593 NACC) is decisive against a coordinate-label permutation null in ADNI (*P <* .001) and, in NACC, is coordinate-localized rather than diffuse: the aggregate AUC only marginally exceeds the null (*P* = .093), but the orientation coordinate carries a clear signal (per-coordinate AUC 0.718; within- versus between-coordinate agreement 91.6% vs 73.5%). These results establish that cognitive decline in AD contains coordinate structure detectable across independent cohorts, instruments (ADAS-Cog and MoCA), and assessment cadences (semi-annual and annual), while also delineating where that structure is and is not recoverable from a given battery.

The ADNI advantage is larger in both absolute log-loss difference and S2 AUC. Two factors likely contribute: (1) ADAS-Cog items span a wider difficulty range and were designed to detect impairment across the MCI-to-dementia spectrum, whereas MoCA sub-items show ceiling effects in cognitively-normal populations; (2) ADNI’s semi-annual assessments provide denser temporal sampling than NACC’s annual visits. The same ceiling mechanism explains the marginal NACC permutation null. Because most MoCA sub-items sit near ceiling and decline together late in disease, the label-shuffled null is high (0.576, versus 0.529 in ADNI): any same-sized grouping of co-declining items reproduces much of the apparent consistency. Coordinate structure becomes recoverable only where items retain discriminative range—the six orientation items, which span a wide severity gradient and show within-coordinate agreement far above the between-coordinate baseline. The honest reading is not that S2 fails in NACC but that the MoCA battery supplies coordinate-specific leverage in only a subset of coordinates; the structural prediction is borne out where the instrument can express it.

### S1 is a sanity check, not the thesis

DCR passes against a pooled continuous baseline in both cohorts, but a mixed-effects IRT model achieves lower log-loss on both (Table 1). This is expected: for *n* patients and *k* items, mixed-effects IRT fits 2*n* + *k* effective parameters against DCR’s *d* coordinate trajectories with a shared deterministic emission, and log-loss is the metric per-patient calibration optimizes. The comparison tells you that mixed-effects IRT is a better probability calibrator at semi-annual or annual cadence. It does not tell you whether cognitive decline has discrete coordinate structure, because the mixed-effects model has no concept of coordinates and cannot produce the S2 equivalence-class signature regardless of parameterization. The thesis is structural: decline is organized by coordinate loss, producing a finite clinical taxonomy rather than smooth drift—a claim that log-loss cannot adjudicate and that no continuous model can produce. The conditions that would falsify this claim are unstructured errors and S2 at chance; cross-task coordinate consistency exceeds chance in both cohorts.

### Relationship to cognitive diagnostic models

The DCR emission is the conjunctive condensation function of the DINA cognitive diagnostic model [13]; the coordinate dictionary is the Q-matrix. We make the correspondence explicit because it both situates DCR in a mature psychometric literature and clarifies what is new. Three differences from classical CDMs matter. First, CDMs are overwhelmingly cross-sectional classifiers of static skill profiles; DCR adds an explicitly longitudinal layer in which attributes are lost over time, connecting to latent Markov [11] and change-point [12] models. Second, the attributes track neurodegenerative dimensions rather than instructional skills, motivating clinical signatures and falsifiable predictions. Third, the *O*(*d*)versus-*O*(2^*d*^) contrast frames DCR’s identifying content against a generic latent-state emission table. We have written the dictionary as a pre-registered Q-matrix so that CDM estimation and validation tools—DINA/DINO variants, higher-order attribute structures, Q-matrix validation— apply directly [14,15].

### Clinical relevance and the S5 null

The S5 null—the inferred mask does not beat the total score for predicting CDR-SB—is consistent across cohorts (*R*^2^ mask 0.497 ADNI, 0.467 NACC vs total 0.674, 0.593). CDR-SB is a global severity measure that collapses across domains, and the total score is already an efficient predictor of global severity. In NACC the 19-item saturated vector substantially exceeds the total (*R*^2^ 0.707 vs 0.593), showing that item-level detail carries clinically relevant information—but the mask’s low-dimensional compression does not capture this advantage. We report S5 as a deliberate, honest null on incremental value for a global outcome, not as evidence against the model. The mask’s potential value lies in predicting domain-specific functional outcomes (navigation impairment from visuospatial loss, medication management errors from executive loss), which CDR-SB does not provide.

The structural claim matters clinically beyond prediction. A coordinate-organized taxonomy (2^*d*^ phenotypes from *d* coordinates) could stratify patients by which capacities are lost, not only how many. With *d* = 5 ADNI coordinates, 32 distinct cognitive phenotypes exist; two patients with identical total scores but different masks would have different functional impairments and potentially different treatment responses. Coordinate-preservation endpoints in trials would detect effects a total score averages away when a drug protects one domain but not others. The model also generates falsifiable predictions about error structure that no continuous model can produce (Supplement S6).

### Encoding is decisive

An earlier NACC analysis using 12 test-level summary scores with z-score binarization and multi-coordinate loading reversed the result (Δ = −0.147); switching to natively discrete sub-items with single-coordinate loading restored it (Δ = +0.036). The reversal was entirely attributable to encoding, not to the data or the model. The product emission operates correctly on discrete single-coordinate probes but produces systematic miscalibration on continuously-scored multi-coordinate aggregates (Supplement S4). This constrains the model’s requirements and implies that earlier CDM applications to dementia data using test-level aggregates would have encountered the same miscalibration.

## Limitations

ADNI assessments occur at approximately 6-month intervals and NACC at annual intervals; the discrete-versus-continuous distinction has maximum leverage at daily-to-weekly sampling and diminishing leverage as interval increases, so change-point dynamics cannot be resolved at either cadence. Single-coordinate encoding makes S3 (ordered vulnerability) and S4 (interaction-order collapse) untestable; testing them requires items with genuine multi-coordinate loading at the item level, not test-level aggregates. The coordinate dictionaries follow standard domain assignments and were not optimized to outcomes; replication across two independently constructed dictionaries reduces but does not eliminate the concern that results depend on a specific battery structure. A sensitivity analysis confirmed the ADNI advantage at all lapse rates and train fractions, but the NACC advantage reverses at *λ* = 0.02 (Δ = −0.045), where the deterministic emission cannot absorb stochastic ceiling-item failures (Supplement S3). Items that are the sole representative of a coordinate are excluded from S2, limiting leverage in single-item coordinates. Abrupt decline can also reflect delirium, depression, medication effects, or vascular events; distinguishing neurodegenerative coordinate loss from reversible state changes remains an open challenge, though reversible dropout-and-recovery is itself informative about which dimensions sit near threshold. The model does not imply abandoning continuous scores; it proposes that some clinically important transitions are lost when only the total is analyzed. Boundary conditions are detailed in Supplement S6.

### Future directions

The discrete-versus-continuous distinction has near-zero statistical leverage at 6-to-12-month sampling, because latent transitions average into a smooth slope, and maximum leverage at the daily-to-weekly sampling that digital assessment now delivers [19–22]. Highfrequency digital cohorts are therefore the decisive arena: item-level data at sufficient temporal resolution would allow resolution of change-point dynamics, dwell times, and recovery after perturbation. Domain-specific functional outcomes (e.g., navigation tasks for visuospatial loss, instrumental activities of daily living for executive loss) would test the mask’s clinical value beyond the total score—the honest predictive bar is improvement over the saturated item-level model, not merely over the total. Cross-disease validation in Parkinson’s cohorts (MoCA items) would test whether coordinate-structured decline is a general property of neurodegeneration rather than AD-specific. Trials can add coordinate-preservation endpoints (delay to loss of a pre-specified discrimination class, reduction in transition probability) alongside CDR-SB or ADAS-Cog.

In summary, the DCR model represents decline as loss of finite discriminative coordinates through a shared-mask constraint with *O*(*d*) identifying content. Its defining signature—cross-task coordinate consistency—replicates across two cohorts, decisively in ADNI and in the orientation coordinate in NACC, while out-of-sample log-loss confirms competitiveness with a pooled baseline but not a per-patient mixed-effects model. The claim is not best-calibrated prediction but that cognitive decline has discrete coordinate structure—requiring natively discrete single-coordinate encoding to detect and that no continuous model can produce. Incremental clinical value for a global outcome was not established; the decisive next arenas are high-frequency digital assessment, domain-specific outcomes, and cross-disease validation.

## Acknowledgments

Data used in preparation of this article were obtained from the Alzheimer’s Disease Neuroimaging Initiative (ADNI) database (adni.loni.usc.edu); ADNI investigators contributed to design and implementation and/or provided data but did not participate in analysis or writing. A complete listing is at https://adni.loni.usc.edu/wp-content/uploads/how_to_apply/ADNI_Acknowledgement_List.pdf. The NACC database is funded by NIA/NIH Grant U24 AG072122, with data contributed by the NIA-funded ADRCs.

## Conflicts of Interest

The author reports no competing interests.

## Funding Sources

None.

## Author Contributions

A.W.: conceptualization, methodology, software, formal analysis, investigation, data curation, writing—original draft, writing—review & editing, visualization.

## Data Availability

ADNI data are available from the LONI Image and Data Archive (https://ida.loni.usc.edu) subject to the ADNI Data Use Agreement; NACC UDS3 data through the NACC data request process (https://naccdata.org). No patient-level data are included in the repository. All code, competing fits, diagnostics, and figure scripts are released for full reproduction; reporting follows STROBE (Supplement).

## Consent Statement

This study is a secondary analysis of de-identified data obtained from the Alzheimer’s Disease Neuroimaging Initiative (ADNI) and the National Alzheimer’s Coordinating Center (NACC) under their respective Data Use Agreements. Per 45 CFR 46, secondary analysis of de-identified datasets does not constitute human-subjects research requiring additional IRB approval. Individual informed consent was obtained by the original data-collecting studies; no additional consent was required for this secondary analysis.

## Notes

https://github.com/aliawu08/dcr-cognitive-resolution

### Competing Interest Statement

The authors have declared no competing interest.

