## Supplementary Material for "Discrete Cognitive Resolution in Alzheimer’s Disease: Cross-Cohort Reanalysis of ADNI and NACC Longitudinal Data"

Alia Wu

**Reproducibility.** All numbers are generated by the released analysis pipeline and figure scripts. Values depending on patient-level data are generated inside the DUA-approved environment; aggregate results are committed in the public repository. Supplementary Note and Table numbers (S1, S2, ...) are referenced from the main text.

#### Supplementary Note S1: The DCR model, full derivation

**State space and deterministic projection emissions.** A cognitive task environment is represented by $d$ binary discriminative coordinates. At time $t$ an individual has an active resolving mask $m(t) = (m_1(t), \ldots, m_d(t)) \in \{0,1\}^d$, where $m_j(t) = 1$ means coordinate $j$ supports reliable discrimination. Resolving power is $\rho(t) = \sum_j m_j(t)$. A stimulus or task demand is a vector $x \in \{0,1\}^d$; the effective record is the projection $R_{m(t)}(x) = m(t) \odot x$. Two states are indistinguishable to the individual when $m(t) \odot x = m(t) \odot y$, so active coordinates partition the environment into $2^\rho$ equivalence classes, each collapsing $2^{d-\rho}$ states distinguishable to a higher-resolution observer. A coordinate-loss event $m_j : 1 \to 0$ halves the distinguishable classes within the engaged subsystem, from $2^\rho$ to $2^{\rho-1}$. The model does not assert literal neural bits; it asserts that clinical tasks decompose into finite discriminative dimensions and that loss of a dimension produces structured coarse-graining rather than uniform noise. The discrete layer concerns the functional consequence once a coordinate falls below the threshold for reliable discrimination, and is compatible with continuous neurobiology (synaptic, network, tau, vascular, inflammatory, sleep, sensory, and medication factors all change the probability that a coordinate remains active).

**The constraint that distinguishes DCR from a generic HMM.** The substantive claim is not that cognition has latent states—mature latent-state and change-point models already encode that—but that emissions are deterministic projections through a shared mask: a task succeeds when all coordinates it requires are active. For task $k$ requiring coordinate set $\tau_k$,

$$P(\text{correct}_k \mid m) = (1 - \lambda)\, \mathbf{1}[\tau_k \subseteq \text{supp}(m)] + \lambda\, \mathbf{1}[\tau_k \nsubseteq \text{supp}(m)],$$

with lapse rate $\lambda$. The emission distribution across all tasks in a state is generated by the single $d$-dimensional mask, not by free per-task parameters. A generic hidden Markov model fits each state's emission vector freely, costing $O(2^d)$ parameters across the state lattice; the DCR constraint ties emissions through the mask at $O(d)$. This restriction is the model's identifying content and the source of its statistical leverage: it predicts that failures across unrelated tasks at a given time are organized by a common set of lost coordinates rather than being task-specific. That cross-task coordinate consistency is the signature that separates DCR from spreading-activation or item-specific confusability accounts, which predict task-local error structure.

**Mask identifiability.** Every clinical use of the model routes through the inferred mask $\hat{m}(t)$. With $d$ coordinates there are $2^d$ possible masks, and from accuracy data alone one cannot in general separate "coordinate inactive" from "coordinate active but failing stochastically" without an externally fixed coordinate dictionary mapping each task to the coordinates it requires. We treat the dictionary as a pre-registered modeling input and mask identifiability as the primary feasibility question. The lapse rate $\lambda$ separates measurement noise from latent state: $P(\text{correct} \mid \text{active}) = 1 - \lambda$, $P(\text{correct} \mid \text{lost}) = \lambda$, and the active-probability inversion is $p_a = (\text{accuracy} - \lambda)/(1 - 2\lambda)$.

#### Supplementary Note S2: Coordinate dictionaries and participant flow

Each item loads on exactly one coordinate (Table S1); single-coordinate loading ensures the product emission reduces to a single active-probability per item. The dictionaries follow the published domain structure of each instrument (MoCA, Nasreddine et al. 2005; ADAS-Cog, Rosen et al. 1984; main-text references [5] and [7]); each sub-item was assigned to the single domain it primarily indexes, and all assignments were fixed before examining outcomes.

In ADNI, the item-level export contained 755 unique participants, of whom 750 had scoreable item data and 688 met the minimum-visit ($\geq 3$) criterion (median 5 visits, IQR 4–5, ∼6-month intervals). In NACC UDS3, 13,323 participants with annual visits met the criterion (of 29,006 with $\geq 3$ visits and 56,532 total). Item-visit observations entering the analyses were 26,022 (ADNI) and 1,089,005 (NACC). Participant flow is shown in Figure S1. The minimum-visit criterion selected a younger, more educated, less impaired analytic sample than the source population (main-text Table 2), as expected when requiring repeated follow-up favors slower-declining participants; findings should be generalized with this selection in mind.

**Table S1:** Coordinate dictionaries (pre-registered Q-matrices). Each item loads on exactly one coordinate.

| Coordinate | ADNI (ADAS-Cog) | NACC (MoCA sub-items) |
|---|---|---|
| Memory | Q1, Q4, Q8 | MOCARECN |
| Language | Q2, Q5, Q10, Q11, Q12 | MOCANAMI, MOCAREPE, MOCAFLUE |
| Visuospatial | Q3 | MOCACUBE, MOCACLOC, MOCACLON |
| Executive | Q6 | MOCATRAI, MOCACLOH, MOCAABST |
| Attention | — | MOCADIGI, MOCALETT, MOCASER7 |
| Orientation | Q7 | MOCAORDT, MOCAORMO, MOCAORYR, MOCAORDY, MOCAORPL, MO[illegible] |

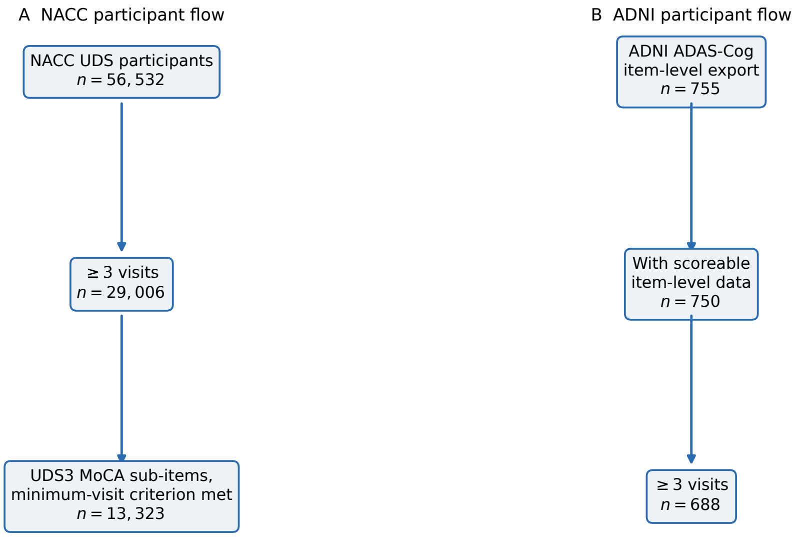


**Figure S1: Participant flow.** (A) NACC: of 56,532 UDS participants, 29,006 had at least three visits; 13,323 met the minimum-visit criterion with UDS3 MoCA sub-items and entered the analysis. (B) ADNI: of 755 participants in the ADAS-Cog item-level export, 750 had scoreable item data and 688 met the minimum-visit ($\geq 3$) criterion and entered the analysis.

### Supplementary Note S3: Sensitivity analysis

The lapse rate ($\lambda = 0.05$) and train fraction (0.66) were fixed before analysis. A pre-specified sensitivity analysis varied $\lambda \in \{0.02, 0.05, 0.10\}$, the train fraction $\{0.50, 0.66, 0.75\}$, and the NACC delayed-recall threshold On ADNI, DCR wins at all lapse rates and train fractions (Table S2). On NACC, DCR wins at $\lambda = 0.05$ and $\lambda = 0.10$ across all train fractions and recall thresholds, but *reverses at* $\lambda = 0.02$ (Table S3): at very low assumed lapse the deterministic product emission cannot absorb stochastic item-level failures on near-ceiling MoCA items ($\Delta = -0.045$ at the primary train fraction). The NACC reversal does not occur on ADNI because the absolute DCR–continuous gap is much larger ($\Delta > 0.25$ even at $\lambda = 0.02$). The result is insensitive to the delayed-recall binarization (Table S4).

**Table S2:** Sensitivity analysis, ADNI ($n = 688$): out-of-sample item-level log-loss. DCR wins at all lapse rates and train fractions.

| $\lambda$ | train frac | Continuous | DCR | Per-item | $\Delta$ (cont−DCR) | Winner |
|---|---|---|---|---|---|---|
| 0.02 | 0.50 | 1.854 | 0.579 | 0.941 | +1.275 | **DCR** |
| 0.02 | 0.66 | 0.965 | 0.490 | 0.833 | +0.475 | **DCR** |
| 0.02 | 0.75 | 0.738 | 0.485 | 0.825 | +0.253 | **DCR** |
| 0.05 | 0.50 | 1.854 | 0.498 | 0.941 | +1.355 | **DCR** |
| 0.05 | 0.66 | 0.965 | 0.436 | 0.833 | +0.529 | **DCR** |
| 0.05 | 0.75 | 0.738 | 0.432 | 0.825 | +0.306 | **DCR** |
| 0.10 | 0.50 | 1.854 | 0.454 | 0.941 | +1.399 | **DCR** |
| 0.10 | 0.66 | 0.965 | 0.410 | 0.833 | +0.555 | **DCR** |
| 0.10 | 0.75 | 0.738 | 0.407 | 0.825 | +0.331 | **DCR** |

**Table S3:** Sensitivity analysis, NACC ($n$ = 13,323): out-of-sample item-level log-loss. DCR wins at $\lambda = 0.05$ and $\lambda = 0.10$ across all train fractions but loses at $\lambda = 0.02$, where the deterministic emission is too rigid to absorb stochastic ceiling-item failures.

| $\lambda$ | train frac | Continuous | DCR | Per-item | $\Delta$ (cont−DCR) | Winner |
|---|---|---|---|---|---|---|
| 0.02 | 0.50 | 0.690 | 0.765 | 0.895 | −0.075 | continuous |
| 0.02 | 0.66 | 0.602 | 0.648 | 0.800 | −0.045 | continuous |
| 0.02 | 0.75 | 0.587 | 0.621 | 0.771 | −0.034 | continuous |
| 0.05 | 0.50 | 0.690 | 0.648 | 0.895 | +0.042 | **DCR** |
| 0.05 | 0.66 | 0.602 | 0.566 | 0.800 | +0.036 | **DCR** |
| 0.05 | 0.75 | 0.587 | 0.547 | 0.771 | +0.040 | **DCR** |
| 0.10 | 0.50 | 0.690 | 0.569 | 0.895 | +0.121 | **DCR** |
| 0.10 | 0.66 | 0.602 | 0.513 | 0.800 | +0.089 | **DCR** |
| 0.10 | 0.75 | 0.587 | 0.500 | 0.771 | +0.088 | **DCR** |

**Table S4:** NACC delayed-recall (MOCARECN) threshold sweep at $\lambda = 0.05$, train fraction 0.66: pass at 2, 3, or 4 of 5 words. DCR wins at every threshold.

| Pass threshold | Continuous | DCR | Per-item | $\Delta$ (cont−DCR) | Winner |
|---|---|---|---|---|---|
| $\geq 2$ of 5 | 0.597 | 0.559 | 0.790 | +0.037 | **DCR** |
| $\geq 3$ of 5 | 0.602 | 0.566 | 0.800 | +0.036 | **DCR** |
| $\geq 4$ of 5 | 0.607 | 0.573 | 0.809 | +0.034 | **DCR** |

#### Supplementary Note S4: The encoding analysis

An earlier NACC analysis used 12 test-level summary scores (MoCA total, Craft Story, MINT naming, digit span, category fluency, Trail Making, Benson figure), z-scored them against a reference group, binarized at $z < -1.5$, and assigned multi-coordinate loadings (e.g. MoCA total $\to$ 5 coordinates). DCR *lost* to continuous ($\Delta = -0.147$). This was entirely an encoding artifact, for two compounding reasons. First, z-scoring and thresholding apply continuous methods to manufacture binary data, discarding the native discreteness the model requires. Second, multi-coordinate product emission on aggregates compounds: with five coordinates each 90% active, $P(\text{pass}) = 0.9^5 = 0.59$, driving predictions toward certain failure even when every coordinate is only mildly impaired. The fix was not to change the model but to use the correct data level—19 natively discrete MoCA sub-items, each loading on a single coordinate, with no z-scoring and no reference group. After the fix, DCR wins on NACC ($\Delta = +0.036$). This constrains the model's requirements: the product emission operates correctly on discrete single-coordinate probes but produces systematic miscalibration on continuously-scored multi-coordinate aggregates. The lesson generalizes—the model is a claim about natively discrete item-level structure, not a transformation to be applied to summary scores.

#### Supplementary Note S5: Cognitive diagnostic models, mixed-effects competitor, and S2 companion analyses

**Relationship to cognitive diagnostic and latent-state models.** The DCR emission is, structurally, the conjunctive ("deterministic-input, noisy-and-gate") condensation function at the heart of the DINA cognitive diagnostic model [S1]: a task succeeds when all required latent attributes are present, modulated by slip and guess parameters that play the role of our lapse rate $\lambda$. The coordinate dictionary is the DINA Q-matrix, and the broader machinery of cognitive diagnostic models [S2] supplies estimation theory, identifiability results, and Q-matrix validation methods that the present minimal implementation does not yet exploit. Three differences matter. First, CDMs are overwhelmingly cross-sectional classifiers of static skill profiles; DCR adds an explicitly longitudinal layer in which attributes (coordinates) are lost over time and the object of inference is the trajectory of the mask, connecting to latent Markov [S3] and change-point [S4] models of cognition. Second, the attributes are hypothesized to track neurodegenerative discriminative dimensions rather than instructional skills, which motivates the clinical signatures. Third, the $O(d)$-versus-$O(2^d)$ contrast is framed against a generic latent-state/HMM emission table, locating DCR's identifying content in the shared-mask restriction. Classical item response theory [S5] is the continuous-drift competitor's home and the appropriate null. We write the dictionary as a pre-registered Q-matrix precisely so that DINA/DINO variants, higher-order attribute structures, and Q-matrix validation tools apply directly.

**Mixed-effects competitor.** Beyond the pooled single-slope continuous-drift baseline, we fit a mixed-effects IRT competitor: item difficulties and a population intercept/time-slope are estimated by pooling all training visits, and each participant receives a random intercept and random time slope shrunk toward the population values by an empirical-Bayes (L2) prior. It achieves lower log-loss than DCR on both cohorts (0.409 vs 0.436 ADNI; 0.380 vs 0.566 NACC; main-text Table 1), as expected for a model with $2n + k$ effective parameters evaluated on calibration's own metric. It has no concept of coordinates and cannot produce the S2 signature.

**S2 companion analyses.** Three pre-specified analyses separate genuine coordinate-specific organization from the global co-decline that any item grouping would capture. (i) *Permutation null*: the coordinate-label vector is shuffled across items (preserving items per coordinate), and the leave-one-item-out S2 AUC is recomputed 1000 times; the P value is the fraction of null AUCs at or above the observed (Table S5). (ii) *Per-coordinate decomposition*: the estimator is stratified by coordinate to localize the signal (Table S6); single-item coordinates (memory in NACC; visuospatial, executive, orientation in ADNI) are excluded for lack of cross-task evidence. (iii) *Coordinate coherence*: within- versus between-coordinate pairwise item agreement (Table S7). The permutation null is the decisive test; the decomposition and coherence analyses are descriptive localizers. In ADNI the observed AUC lies $\approx 5$ SD above the null; in NACC the null is high (random groupings of co-declining, ceiling-prone items reproduce much of the apparent consistency), leaving the aggregate marginal while the orientation coordinate carries a clear signal.

**Table S5:** S2 permutation null (seed 20260610). The P value is the fraction of 1000 label-shuffled null AUCs $\geq$ the observed AUC.

| Cohort | Observed AUC | Null mean | Null SD | Null 95% CI | $P$ | $n_{\text{perm}}$ |
|---|---|---|---|---|---|---|
| ADNI ($n = 688$) | 0.782 | 0.529 | 0.048 | [0.46, 0.66] | $< 0.001$ | 1000 |
| NACC ($n =$ 13,323) | 0.593 | 0.576 | 0.013 | [0.56, 0.60] | 0.093 | 1000 |

**Table S6:** Per-coordinate S2 decomposition. Coordinates ordered by AUC within cohort; single-item coordinates excluded. The NACC signal concentrates in orientation—the coordinate with the widest discriminative range and least ceiling compression.

| Cohort | Coordinate | Items | AUC | Err. (weak) | Err. (intact) | Item-visits |
|---|---|---|---|---|---|---|
| ADNI | memory | 3 | 0.673 | 0.86 | 0.60 | 9,818 |
| | language | 5 | 0.614 | 0.61 | 0.06 | 16,204 |
| NACC | orientation | 6 | 0.718 | 0.64 | 0.04 | 397,820 |
| | executive | 3 | 0.599 | 0.60 | 0.22 | 178,193 |
| | language | 3 | 0.567 | 0.49 | 0.24 | 191,993 |
| | attention | 3 | 0.547 | 0.27 | 0.16 | 142,786 |
| | visuospatial | 3 | 0.514 | 0.30 | 0.18 | 178,213 |

**Table S7:** Coordinate coherence. "Agreement" means two co-observed items both pass or both fail. Items sharing a coordinate agree more than items on different coordinates in both cohorts; the NACC excess is carried by orientation (within-coordinate agreement 91.6%).

| Cohort | Within-coord | Between-coord | Excess | Within / between pairs |
|---|---|---|---|---|
| ADNI ($n = 688$) | 85.3% | 51.2% | +34.1 pp | 42,114 / 135,833 |
| NACC ($n = 13{,}323$) | 83.2% | 73.5% | +9.7 pp | 1,622,965 / 8,097,314 |

Per-coordinate within-coordinate agreement. **ADNI:** language 90.6%, memory 67.7%. **NACC:** orientation 91.6%, attention 73.9%, visuospatial 70.5%, executive 70.1%, language 67.2%.

#### Supplementary Note S6: Falsifiable predictions and boundary conditions

**Falsifiable predictions.** The DCR model generates five pre-specified falsifiable predictions. Each is stated with its current evidentiary status.

1. Trajectories at sufficient temporal frequency show latent change points that improve out-of-sample item-level prediction over smooth drift. *(Supported over the pooled continuous baseline in both cohorts: ADNI* $\Delta = 0.529$*, 95% CI [0.44, 0.63]; NACC* $\Delta = 0.036$*, 95% CI [0.029, 0.043]. A mixed-effects IRT model achieves lower log-loss than DCR on both cohorts; partially supported—DCR beats the pre-specified pooled baseline but not a per-patient random-effects model optimized for the same metric.)*

2. Error distributions show equivalence-class structure, consistently across unrelated tasks. *(LOO AUC 0.782 ADNI, 0.593 NACC. Decisive against the coordinate-label permutation null in ADNI (*$P < .001$*); in NACC the aggregate is marginal (*$P = .093$*) but the signal concentrates in orientation (per-coordinate AUC 0.718), the coordinate least affected by ceiling.)*

3. Multi-coordinate tasks lose performance earlier than single-coordinate tasks after difficulty adjustment. *(Not testable with single-coordinate item encoding.)*

4. Interaction-order (Walsh–Hadamard [S6]) decompositions show loss of high-order terms before low-order terms. *(Not testable on ADAS-Cog or MoCA; requires three-coordinate tasks with binary sub-items.)*

5. Patients with similar total scores but different inferred masks show different functional outcomes. *(Not established; S5 null on increment over total for CDR-SB in both cohorts. Requires domain-specific functional outcome measures.)*

**Boundary conditions.** The DCR model has identifiable boundary conditions that constrain its applicability and should guide future applications.

*Lapse rate sensitivity.* The lapse rate $\lambda$ governs how rigidly the deterministic emission absorbs stochastic item-level noise. At $\lambda = 0.02$, each misprediction on an "active" coordinate costs $-\log(0.02) = 3.91$ nats. MoCA ceiling items (6 of 19 with mean accuracy $> 0.9$) produce stochastic failures the model cannot absorb at low lapse, reversing the NACC advantage ($\Delta = -0.045$ at $\lambda = 0.02$; Supplementary Table S2). The ADNI advantage is robust across all lapse rates because the absolute DCR–continuous gap is much larger ($\Delta > 0.25$ even at $\lambda = 0.02$). The boundary is instrument-specific: batteries with many near-ceiling items are more sensitive to lapse-rate specification.

*Single-coordinate encoding constraint.* The present analyses use single-coordinate item loading exclusively, which makes S3 (ordered vulnerability) and S4 (interaction-order collapse) untestable. Testing these signatures requires items with genuine multi-coordinate loading at the item level—for example, complex reasoning tasks that simultaneously require executive function and visuospatial processing. This is a constraint of the available item-level data, not of the model itself.

*Assessment cadence.* The discrete-versus-continuous distinction has maximum leverage at daily-to-weekly sampling, where coordinate transitions produce detectable step changes in item-level accuracy. At semi-annual (ADNI) and annual (NACC) cadence, transitions that occur between assessments are averaged into apparent smooth slopes, limiting the model's ability to detect change-point dynamics. High-frequency digital cognitive assessment is the decisive arena for testing the model's temporal predictions.

*Global versus domain-specific outcomes.* The S5 null—the inferred mask does not beat the total score for predicting CDR-SB—is expected for a global severity measure that the total already captures efficiently. The mask's $d$-dimensional compression is too coarse to compete with the total for a global outcome and too coarse to capture the item-level detail that the saturated vector exploits. The mask's potential clinical value would appear for domain-specific functional outcomes (navigation impairment from visuospatial loss, medication management errors from executive loss), which CDR-SB does not provide.

*Confounds with reversible state changes.* Abrupt decline in item-level performance can reflect delirium, depression, medication effects, or vascular events rather than permanent neurodegenerative coordinate loss. The model does not distinguish irreversible coordinate loss from reversible state changes; longitudinal designs with sufficient temporal resolution and clinical adjudication are needed to separate these.

### STROBE checklist

Reporting follows the STROBE guideline for observational studies; the completed item-by-item checklist is provided in the repository.

[S2] Rupp AA, Templin J, Henson RA. Diagnostic Measurement: Theory, Methods, and Applications. New York: Guilford Press; 2010.

[S3] Bartolucci F, Farcomeni A, Pennoni F. Latent Markov Models for Longitudinal Data. Boca Raton, FL: CRC Press; 2013.

[S4] Hall CB, Ying J, Kuo L, Lipton RB. Bayesian and profile likelihood change point methods for modeling cognitive function over time. Comput Stat Data Anal. 2003;42:91-109.

[S5] Embretson SE, Reise SP. Item Response Theory for Psychologists. Mahwah, NJ: Lawrence Erlbaum; 2000.

[S6] Walsh JL. A closed set of normal orthogonal functions. Am J Math. 1923;45:5-24.
